# A self-aligning end-effector robot for individual joint training of the human arm

**DOI:** 10.1101/2020.07.09.195073

**Authors:** Sivakumar Balasubramanian, Sandeep Guguloth, Javeed Shaikh Mohammed, S. Sujatha

**Affiliations:** Department of Bioengineering, Christian Medical College, Vellore, Tamil Nadu 632002, India; Department of Mechanical Engineering, Indian Institute of Technology - Madras, Chennai, Tamil Nadu 600036, India

**Keywords:** arm, rehabilitation, robotics, mechanisms, ergonomics, self-aligning

## Abstract

Current evidence indicates that individual joint training with robotic devices can be as effective as multi-joint training for the arm. This makes a case for developing simpler and more compact robots for training individual joints of the arm. Such robots have the highest potential for clinical translation. To this end, the current work presents the kinematic design and optimization of a six degrees-of-freedom (*dof*) end-effector robot with three actuated *dof* and three non-actuated self-aligning *dof* for safe assisted training of the individual joints (shoulder or elbow) of the human arm, with relaxed constraints of the relative positioning of the human limb with respect to the robot. Further, we present a simple estimation procedure to automatically identify the kinematic parameters of the human limb essential for control of the human-robot closed kinematic chain.

## 1 Introduction

Robot-assisted rehabilitation of arm function, in chronic stroke patients, has been found to be as effective as intensity matched conventional therapy for reducing impairments, but there appears to be little or no effect on functional ability [1, 2, 3]. A study by Milot et al. used the BONES robot to administer multi-joint and single-joint training in moderately impaired chronic stroke subjects, and found that both groups showed improvements in both motor function and impairments [4]. Interestingly, they did not find the multi-joint training to be superior to the single-joint training approach. Although only a pilot study with 20 subjects, this finding is supported by other evidence in the literature. Schaefer et al. found that training on a single activity of daily living (ADL) generalized to untrained tasks in chronic stroke subjects [5]. The study by Fluet et al. investigating robot-assisted upper limb (UL) training showed that training the arm and hand together as a functional unit was no different from training the arm and hand separately [6]. A study on healthy individuals by Klein et al. found that individual training of parts of a complex movement resulted in slightly better learning and retention [7]. From the perspective of developing robots for rehabilitation, these results support the idea of developing simpler, compact, and cost-effective robots that can be used for training one or two degrees-of-freedom (*dof*) at individual joints, rather than developing a single robot capable of training highly complex multi-joint movements. This idea is the primary objective of the current work, which focuses on the development of a robot-assisted training of two *dof* of the shoulder or elbow joint.

*Exoskeleton* robots are well suited for supporting and assisting movements of individual joints [8, 9, 10]. However, exoskeleton robots require close alignment of the human and robot joint axes to ensure safe physical human-robot interaction (pHRI). Misalignment in orientations and offsets between the different human and robot joint axes can result in unwanted forces and torques on the human limb, which means that attaching a human limb to the robot requires care. Furthermore, the robot’s exact kinematic structure is also highly dependent on the topology of the human joint’s kinematic structure. On the other hand, *end-effector* robots have kinematic structures that are independent of that of the limb [11, 12, 13, 14]. They also have significantly fewer constraints on the location and the orientation of the human joint with respect to the robot, thus making it much easier to interface a limb with the robot. Given these advantages, we chose to implement an end-effector based approach for the proposed robot for training individual joints.

Conventional end-effector robots for arm rehabilitation are attached to the human limb at the hand, where they apply interaction forces to impose arm movements. Such an approach cannot be used to impose movements at an individual joint of the arm as this human-robot closed kinematic chain is under-constrained. An end-effector robot for training a specific joint will need to be attached to part of the human limb anatomically connected to the joint of interest (e.g., upper-arm for the shoulder and forearm for the elbow). In order to ensure that the robot imposes precise movements and applies safe forces/torques on the human limb, it is essential for the robot to be aware of the details of the human limb’s kinematic chain and its parameters. Current end-effector robots do not take into consideration the details of the human limb’s kinematic chain.

To this end, we present a 6-*dof* end-effector robot AREBO (**A**rm **Re**habilitation Ro**bo**t) with three actuated (active) *dof* and three unactuated (passive) *dof*. The current paper focuses on the details of AREBO’s kinematic chain, its optimization, and an algorithm for continuously tracking the parameters of the human limb’s kinematic chain. AREBO can support and assist two *dof* of a human joint, which we assume as the human shoulder joint for demonstration purposes. We also present a simple approach for automatically estimating and tracking the human limb’s kinematic chain parameter, which will be used to control the robot-human closed kinematic chain.

## 2 Design of AREBO’s Kinematics Chain

The objective was to develop a compact, portable robot for training movements of individual joints of the human arm (shoulder and elbow), which can be used for both the left and right arms without requiring any change to the robot’s structure. Furthermore, we also wanted to avoid the need for precise positioning and orientation of the patient with respect to the robot, which can be difficult and time consuming with severely affected patients. These design requirements can be fulfilled by an end-effector type robot, the type chosen for designing AREBO.

Consider the human limb with a joint (e.g., shoulder joint) located at the origin of a reference frame 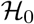 depicted in Fig. 1 with a rigid body (e.g., upper-arm depicted as a red ellipse) with a reference frame 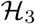 attached to it. This rigid body can undergo pure rotational movements with respect to the frame 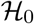. Let 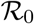 be an earth-fixed reference frame, which also acts as the base of the robot that interfaces to the human limb. We assume that the robot’s endpoint is attached rigidly to the origin of 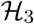, which is the point of physical interaction between the robot and the human. The homogenous transformation matrices 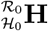 and 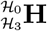 represent the frame 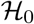 with respect to 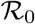, and frame 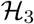 with respect to 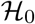, respectively.

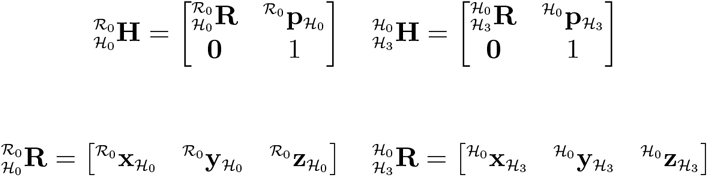

where, 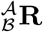 is the rotation matrices representing frame 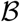 with respect to 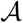, and 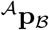 is the location of the origin of frame 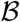 represented with respect to the origin of frame 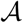.

**Figure 1:**
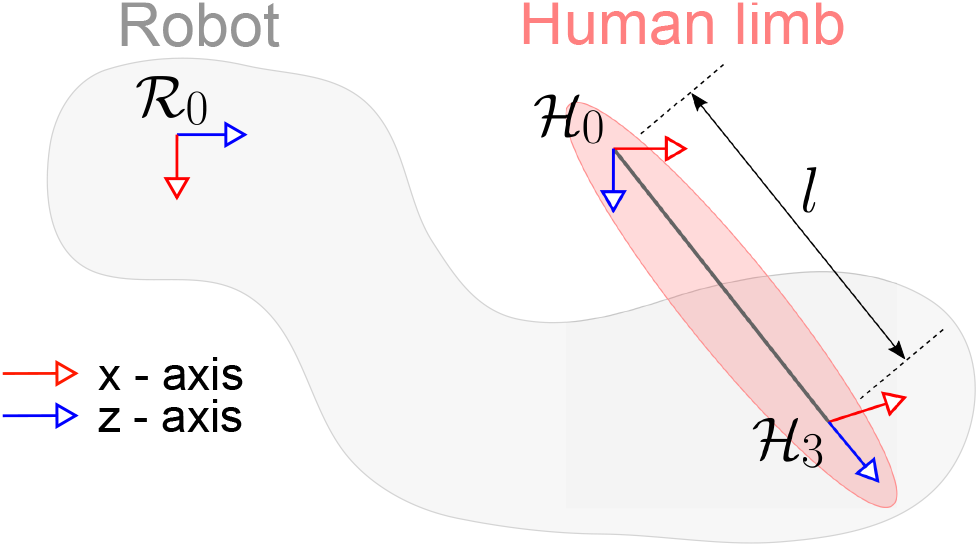
Depiction of a human-robot closed-loop kinematic chain where the movements of the human limb are to be assisted by the robot.

To simplify the process of connecting the robot to a human limb, the robot must not strictly constrain the location and orientation of the human limb with respect to the robot, i.e., no strict constraints on 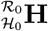. Additionally, the robot must also accommodate human limbs of different sizes, i.e., variations in *l* between different subjects (Fig. 1).

### 2.1 Human Limb’s Kinematic Chain

We assume the human shoulder joint at 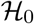 to be a spherical joint realized as three intersecting orthogonal revolute joints (Fig. 2) with generalized coordinates ****ϕ**** (*t*) = [*ϕ*_1_ (*t*) *ϕ*_2_ (*t*) *ϕ*_3_ (*t*)]^*T*^, where *ϕ*_1_ (*t*) and *ϕ*_2_ (*t*) are the shoulder flexion/extension (SFE) and shoulder abduction/adduction (SAA) angles, respectively, at time *t*.

**Figure 2:**
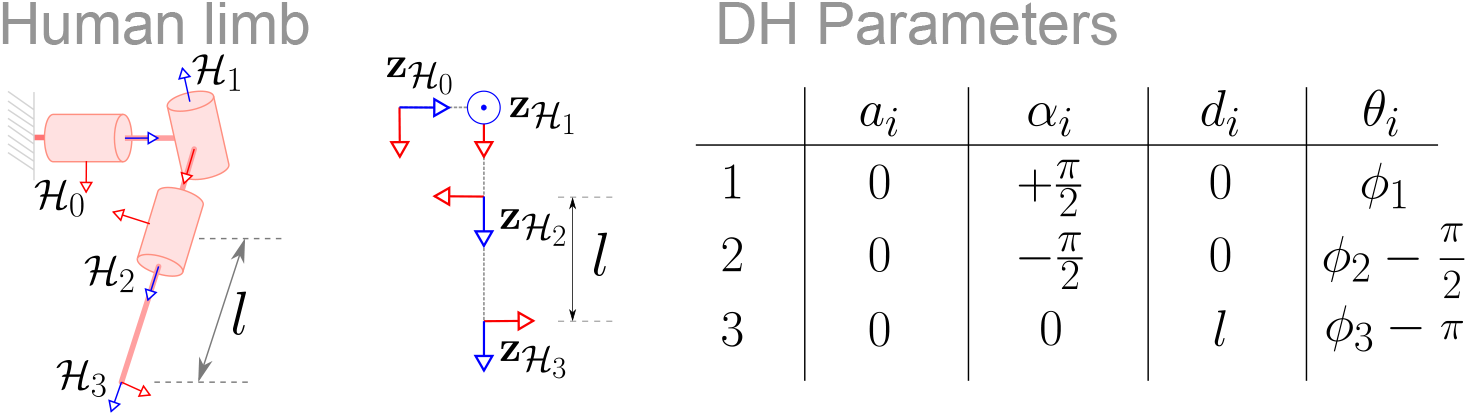
Details of the human limb’s kinematic chain. The human limb considered in this work is a two or three *dof* chain with the structure shown in the figure. The third *dof* is optional.

The DH parameters for the human limb are shown in Fig. 2. The position and orientation of 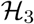 with respect to 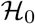 is given by,

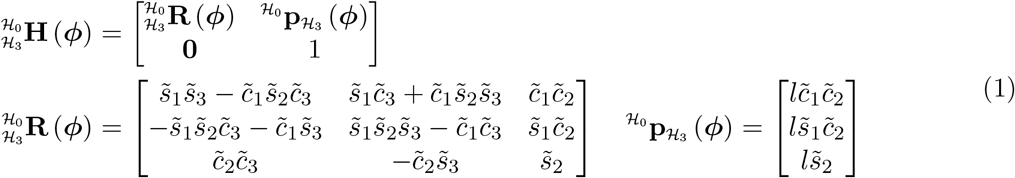

where, 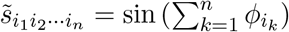 and 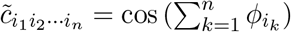.

### 2.2 AREBO’s Kinematic Chain

We are only interested in assisting the SFE and SAA movements of the shoulder joint with the robot in the current application. This movement assistance can be accomplished by applying forces orthogonal to 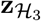, which will result in pure moments about 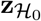 and 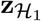, and ensure there are no forces along the length of the human limb that push it into or pull it away from the shoulder joint [15]. This feature requires the robot to possess the following capabilities:

1. It must **apply forces in any arbitrary direction** in space with respect to 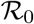. Thus, ensuring it can apply forces orthogonal to 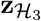, which can have an arbitrary direction depending on the location of the shoulder joint and then the joint configuration of the human limb.
2. The robot needs precise **measurement of the orientation of** 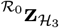 to be able to apply forces orthogonal to 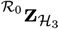.
3. The robot must **align to any arbitrary orientation of** 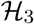 with respect to 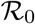, to prevent applying unwanted moments at the interface between the human limb and the robot.

An appropriately designed 6-*dof* robot can achieve any arbitrary position and orientation within its workspace. To apply a force in any arbitrary direction at the robot’s endpoint in 3D space, we need at least three actuated *dof* for the robot. The remaining three *dof* can be unactuated, allowing them to self-align to any arbitrary orientation of the human limb 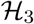. A schematic of the current design for AREBO’s kinematic chain is shown in Fig. 3, which has 6 revolute joints arranged in the specific order shown in the figure. The robot’s generalized coordinates are represented by ***θ*** (*t*) = [*θ*_1_ (*t*) *θ*_2_ (*t*) *… θ*_6_ (*t*)] ^*T*^, where *θ_i_* (*t*) *∈* [0, 2*π*) is the generalized coordinate corresponding to the *i^th^* revolute joint in Fig. 3. The position and orientation of the robot’s endpoint frame 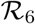, represented by 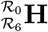 depends on ****θ**** (*t*). The DH parameters of AREBO’s proposed kinematic structure is also listed in a table in Fig. 3; the three parameters *r*_1_, *r*_2_, and *r*_3_ are the different robot link lengths. The resulting homogenous transformation matrix representing 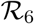 in 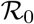 is as follows,

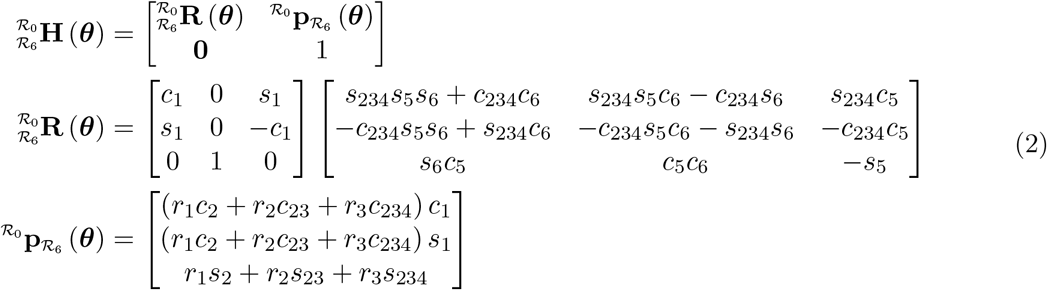

where, 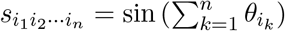 and 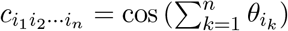.

**Figure 3:**
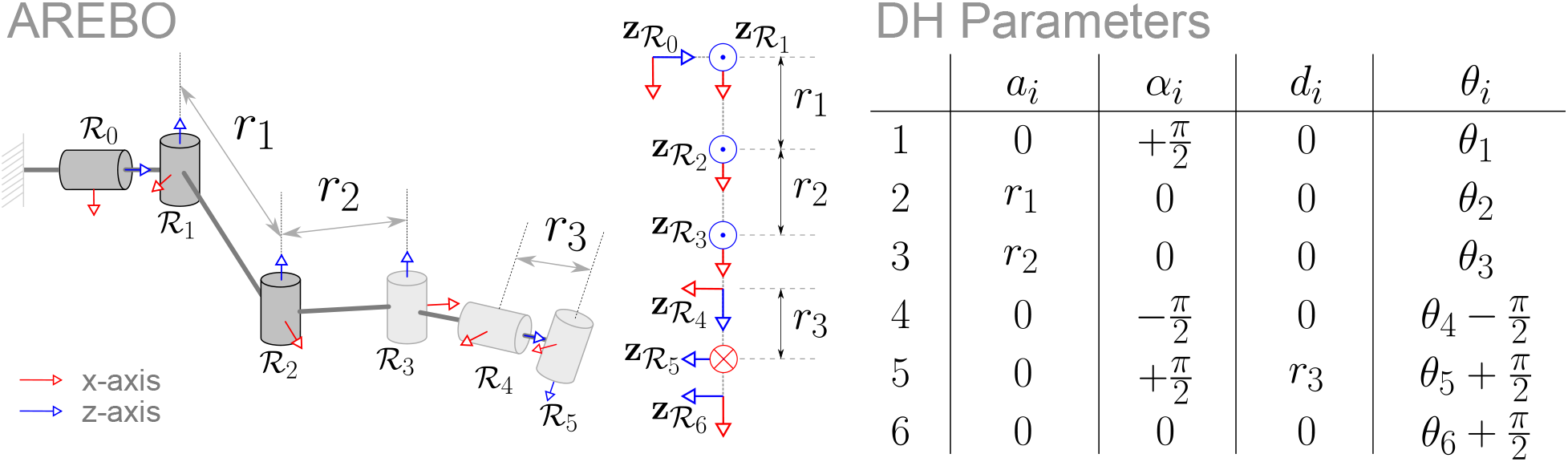
Details of the proposed robot’s kinematic chain. The robot has 6 *dof* arranged in the particular manner shown in the figure. This allows the robot to achieve a range of positions and orientations within its reachable workspace. The first three *dof* (shown in dark gray) are actuated, while the rest three *dof* (shown in light gray) are passive self-aligning joints.

### 2.3 Human-Robot Closed Kinematic Chain

A closed-loop kinematic chain is formed when AREBO is connected to the human limb, such that the frames 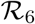 and 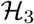 match in position and orientation.

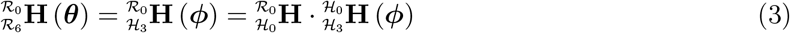

We assume that the orientation of 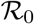 and 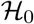 are the same, but they are displaced, i.e.,

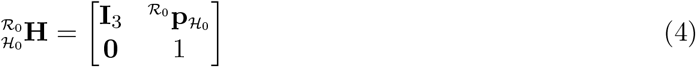

where, **I**_3_ is a 3 *×* 3 identity matrix.

The first three joints of the robot shown in a darker color in Fig. 4 are the actuated joints, while the rest three (shown in a lighter shade) are unactuated. These sections are shown separately in Fig. 4 to demonstrate the different roles of the two sections. The actuated section is responsible for applying appropriate forces on the human limb, while the unactuated section helps in self-aligning the robot’s endpoint 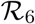 to the human arm 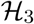. The force resulting from the application of torques *τ*_1_*, τ*_2_*, τ*_3_ at the three actuated robot joints will be transmitted to the human limb through the robot’s unactuated section. In order to induce pure moments about the first two joints (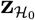 and 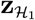 in Fig. 4) of the human limb, the robot’s endpoint force 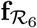 must be orthogonal to 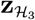.

**Figure 4:**
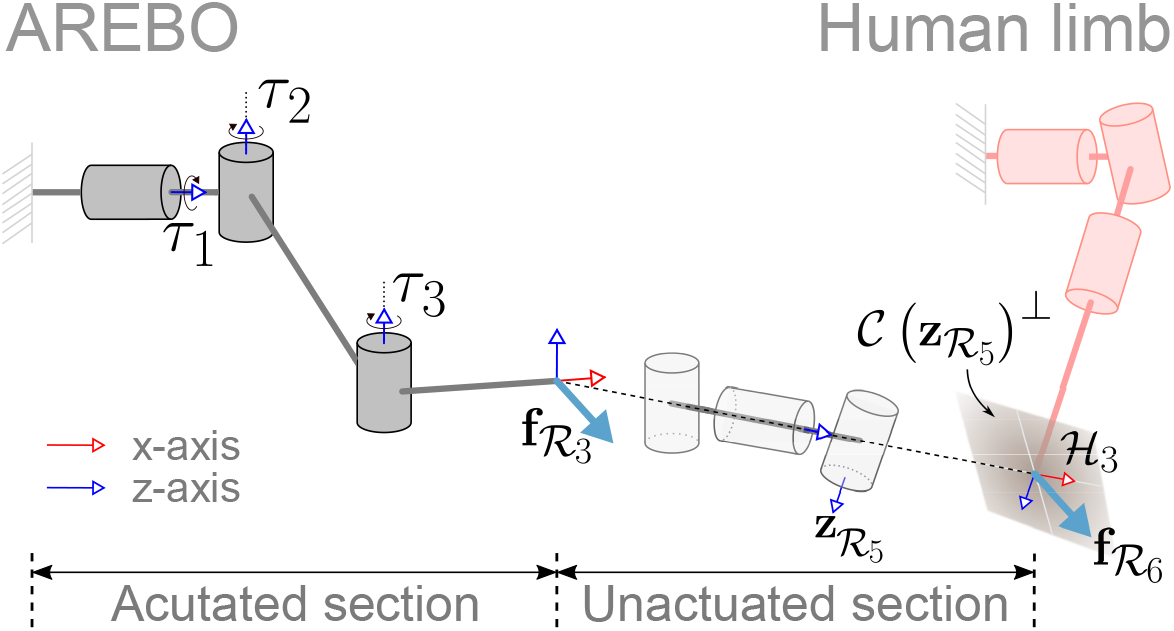
Detailed depiction of the human-robot closed kinematic chain, along with the interaction force applied by the robot on the human limb. The endpoint force 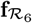 on the human limb is determined by the torques acting on the first three actuated *dof*, which need to be appropriately chosen to ensure 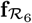 is orthogonal to 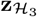.

When the robot and the human limb are connected (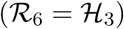), the robot’s 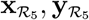 are orthogonal to the human limb’s 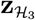 of the human. Applying torques at the three actuated joints 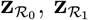, and 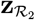 will result in a force 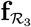 generated at the origin of 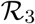 (Fig. 4).

The force 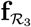 generated at the origin of 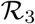 by applying torques at 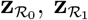, and 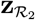 is transmitted to the human limb through the robot’s unactuated section, i.e., 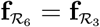, which we would like to be orthogonal to 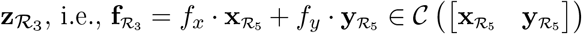, where *f_x_, f_y_* ∈ ℝ. The relationship between 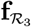 and the torque ****τ**** = [*τ*_1_ *τ*_2_ *τ*_3_]^*T*^ is given by the following relationship,

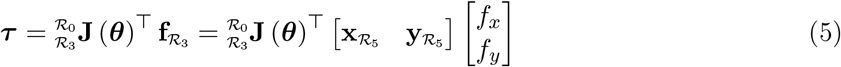

where, 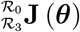 is the Jacobian matrix relating the angular velocities 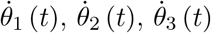 to the linear velocity of the origin of 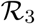. The above equation can be solved for ****τ**** to apply any force 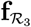 that is in the column space of 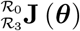. The Jacobian matrix, 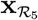, and 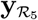 are given by the following,

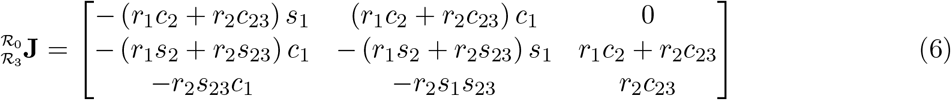

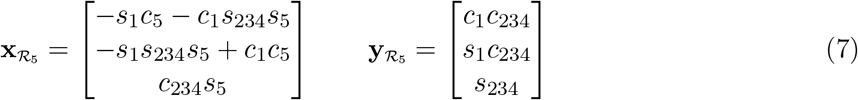

Except for the case where 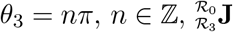 is always full rank and thus any 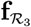 can be applied by appropriately choosing ****τ****.

## 3 Identification of Human Limb Parameters

In the human-robot closed kinematic chain, the planning and control of the human limb’s movements require information about the location of 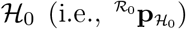 and the length of the human limb *l*. Knowledge of these parameters is crucial for answering the following two specific questions:

1. Can a given desired human limb configuration (*ϕ*_1_*, ϕ*_2_) be reached by the human-robot closed kinematic chain, i.e., *∃**θ***, 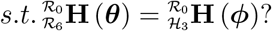
2. For any reachable human limb configuration (*ϕ*_1_*, ϕ*_2_), what is the corresponding robot configuration ****θ**** that allows the human limb to achieve (*ϕ*_1_*, ϕ*_2_)?

Assuming that we know 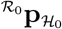 and *l*, we can answer the two questions by first performing forward kinematics for the human limb to compute 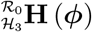 for a given (*ϕ*_1_*, ϕ*_2_); we assume *ϕ*_3_ to be anything as any rotation about this *dof* is accommodated by *θ*_6_. We then perform inverse kinematics for the robot by assuming 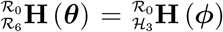. The algorithm for performing inverse kinematics for AREBO is detailed in Appendix A. If we obtain a valid ****θ**** for a given (*ϕ*_1_*, ϕ*_2_), then this human limb configuration is achievable and ****θ**** is the corresponding robot joint configuration.

Given that AREBO allows some freedom for the user to sit with respect to the robot (Fig. 7) and can accommodate upper-limbs of different sizes, the values of 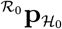 and *l* will be different for different users. It is not practical to measure these parameters for each user in order to use the robot. To address this issue, we propose a simple calibration procedure that can automatically estimate these parameters once a user is connected to the robot. This calibration can be done through a least squares estimation procedure, where the robot imposes random, safe movements to the human limb while recording the ****θ**** of the robot, and the *pitch* (*ϕ*_1_) and *yaw* (*ϕ*_2_) angles of the human limb. Let us assume that we have a record of the robot and human limb angles.

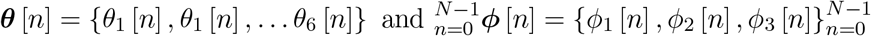

where, *n ∈* ℤ is the time index, and *N ∈* ℤ, *N >* 0 is the length of data available. In the human-robot closed kinematic chain, we have

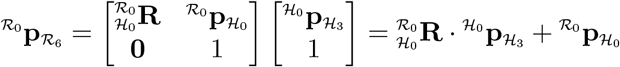

**Figure 7:**
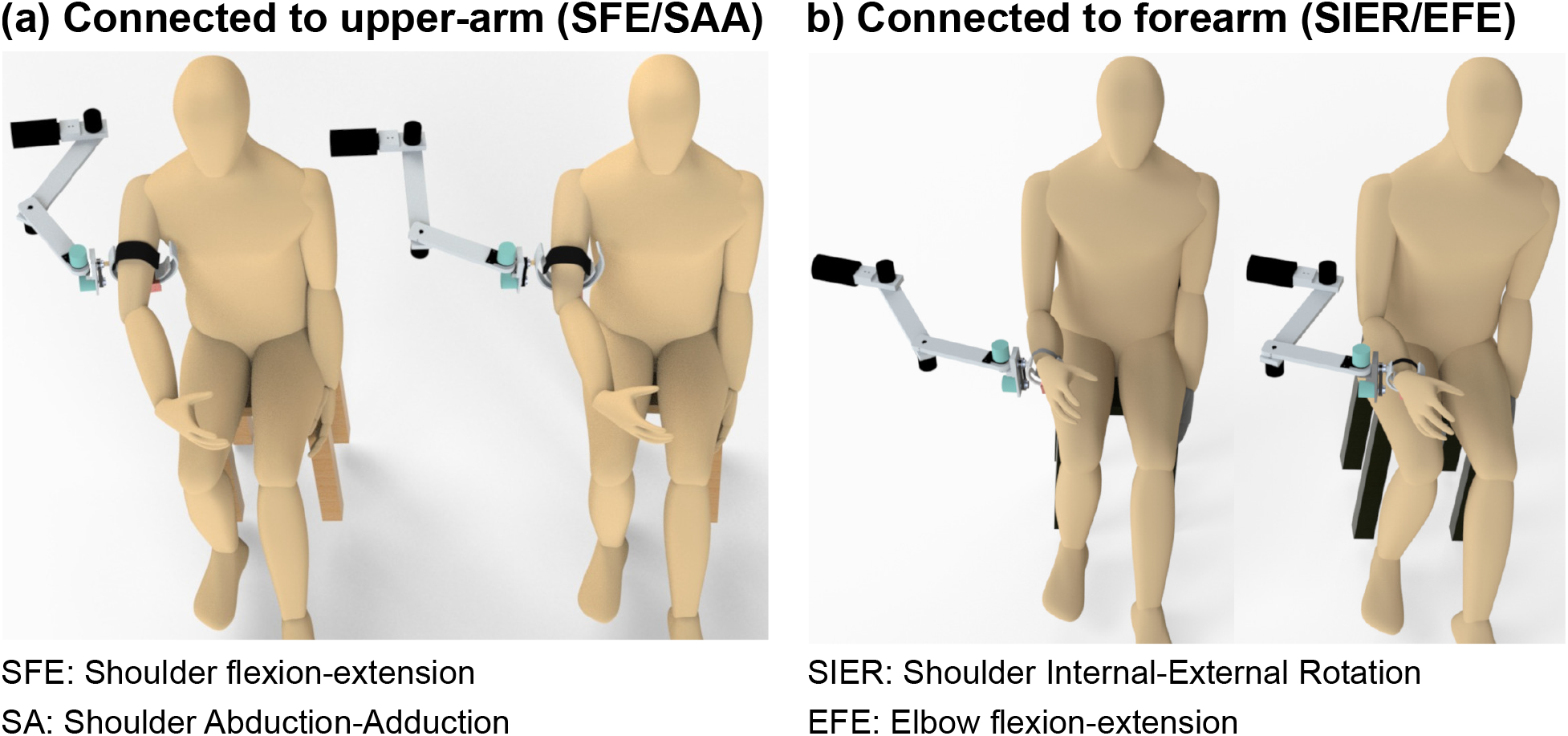
CAD models of a realization of AREBO and the depiction of its use for assisting the different movements are individual joints. (a) When connected to the upper-arm, the robot can assist SFE and SAA by applying forces orthogonal to the upper-arm. (b) When it is connected to the forearm, SIER and EFE can be assisted by applying forces orthogonal to the forearm. It should be noted that this approach for assisting SIER is safer and more comfortable than by providing tangential forces on the upper-arm [15]. The two CAD models in (a) and (b) demonstrates the relative freedom a subject has in sitting with respect to the robot.

We have assumed earlier that 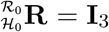, and let 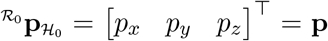. Then, for any time instant *n* we have from the above equation, Eq. 2, and Eq. 1,

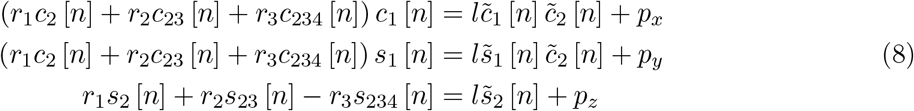

The unknowns in the above equation are *p_x_, p_y_, p_z_*, and *l*. We can rewrite the above equation in the following form,

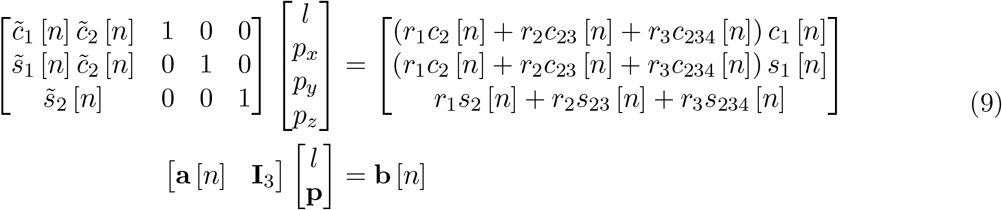

Combining the equations for all *n*

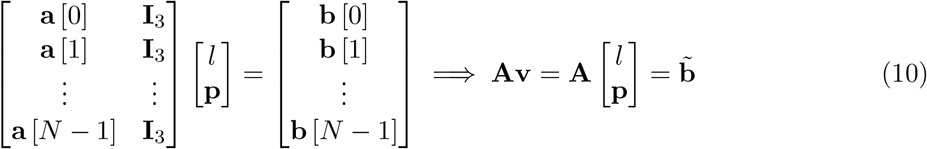

If **A** is full rank, the least squares estimate of the parameters are given by,

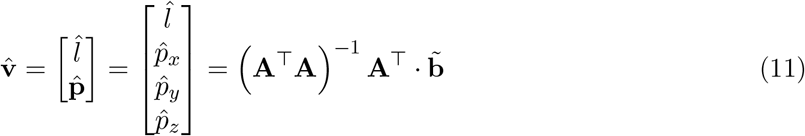

## 4 Methods

In this section, we describe the optimization of the robot link length parameters and the simulation analysis of the algorithm for estimating human limb parameters.

The first step in the physical realization of AREBO is the choice of its link lengths *r*_1_, *r*_2_, and *r*_3_. These link lengths will determine the robot’s workspace and its overall manipulability. The individual endpoint workspaces of the robot 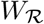 and the human limb 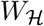 are given by the following,

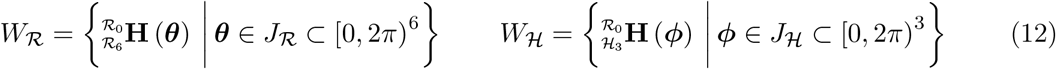

where, 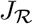 and 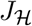 are the set of all joint configurations that can be achieved by the robot and the human limb, respectively. 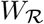 and 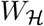 are the set of all endpoint positions and orientations of the robot and the human limb. Here, all positions and orientations are represented with respect to the common frame 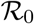.

When the endpoints of the human limb and robot are attached together, i.e., 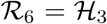, the only possible joint configurations 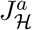 and 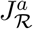 for the human limb and the robot are those corresponding to the set 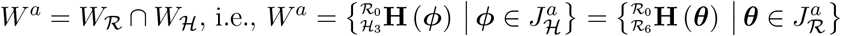. In general there will be a reduction in the human limb’s workspace 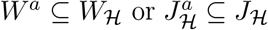. The size of the set *W*^*a*^ is determined by several factors: (i) length of the robot links *r*_1_*, r*_2_, and *r*_3_; (ii) length of the human limb *l*; and (iii) position of the human limb with respect to the robot 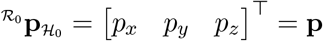.

We wish to allow a range of possible values for the parameters *l* and **p** to support the movements of human limbs of different lengths without placing strict constraints on the seating of a subject with respect to the robot. Thus, for given reasonable parameter ranges for *l* and **p**, we would like to choose the robot link lengths to achieve two objectives:

1. Maximize *W^a^* (or 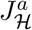) to minimize the restrictions on the human limb’s movement, and
2. Maximize the ability of the robot to apply forces orthogonal to the human limb.

These two objectives can be combined into a single objective function *O* (**r**) of the robot link length **r** = [*r*_1_ *r*_2_ *r*_3_]^*T*^,

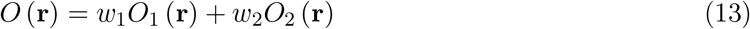

where, *O*_1_ (**r**) is a function of *W^a^*, and *O*_2_ (**r**) is a function of the robot’s ability to apply forces in the appropriate directions, 0 *< w*_1_*, w*_2_ ∈ ℝ are the weights for the two objectives. It should be noted that *O*_1_ (·) and *O*_2_ (·) are only functions of **r** as these are obtained by averaging these measures across the different possible values of *l* and **p**. The optimal value for the robot’s link lengths **r**^*opt*^ is obtained by maximizing *O* (**r**) for a given range of values for *l* and **p**.

### Human limb’s workspace *O*_1_ (r)

The first objective function *O*_1_ (**r**) depends on the human limb’s workspace when it is connected to a robot with link lengths **r**. We chose to quantify the size of the workspace in the joint space 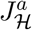, where 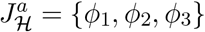. For a given value of **r**, *l*, and **p**, we quantify the normalized workspace of the human limb as the following,

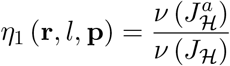

where, *ν* (·) computes the volume of a given set, which implies that 0 ≤ *η*_1_ (**r***, l*, **p**) ≤ 1. *O*_1_ (**r**) is obtained for a given value of **r** by computing the average value of *η*_1_ (**r***, l*, **p**) over the range of values for *l* and **p**,

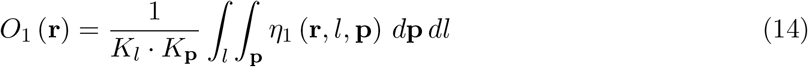

where, *K_l_* = *∫l dl* and *K*_**p**_= **∫p***d***p** are the sizes of sets of parameters *l* and **p**, respectively.

### Robot manipulability *O*_2_ (r)

Manipulability provides a measure of how easily a robot can apply forces in different directions, which, in general, depends on the robot’s joint configuration. In the current application, we are not interested in applying forces in any direction, but only in the plane orthogonal to the human limb. In the closed kinematic chain shown in Fig. 4, for a given **r**, *l*, and **p**, for any human limb configuration in 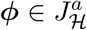, there is a corresponding point 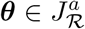. The relationship between the torque ****τ**** at the robot’s joints and its endpoint force 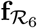 depends on the robot’s Jacobian matrix for the current joint configuration ****θ**** (Eq. 5). For any given torque the resulting force along the human limb (**f**_*z*_) and orthogonal to the human limb (**f**_*xy*_) can be obtained from the following expressions,

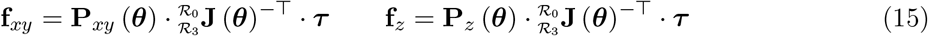

where, 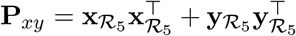 and 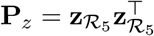 are the orthogonal projection matrices on to the *xy*-plane and the *z* axis of the frame 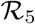. We would ideally like the robot’s kinematic chain to be inherently more suited to apply forces in the *xy*-plane of 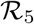, rather than the *z*-axis, for a given ****τ****. This property is captured by the following measure, which captures the ratio of the maximum possible force in the *xy*-plane with respect to that of the *z*-axis for a given torque ****τ****.

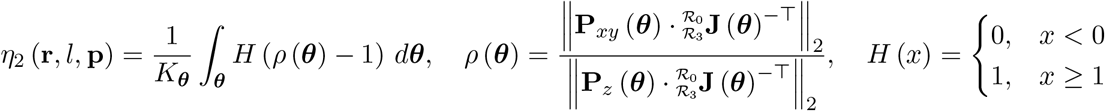

where, *K_**θ**_* = ****∫_θ_*** d***θ**** is the size of 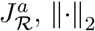 is the induced second norm of a given matrix, and *H* (*·*) is the step function. The ratio *ρ* (****θ****) is a measure of the “ease” of applying a force along the *xy*-plane compared to that of *z*-axis at the joint configuration ****θ****. Ideally, we would like this ratio to be greater than 1, and thus this ratio is transformed using the step function, such that ratios that are less than 1 are mapped to 0, and the ones greater than or equal to 1 are mapped to 1. Thus, *η*_2_ (**r***, l*, **p**) can be interpreted as the proportion of points in the robot’s workspace with the ratio *ρ* (****θ****) *≥* 1, which implies that 0 *≤ η*_2_ *≤* 1.

*O*_2_ (**r**) is obtained from *η*_2_ (*·*) by averaging over the other two arguments *l*, and **p**(like Eq. 14).

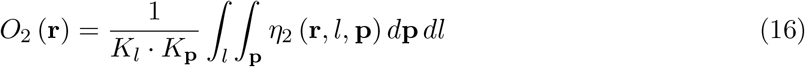

#### 4.1 Optimization of robot link lengths

The optimization of the robot link lengths was carried out numerically through a brute force search over a set of parameters values for **r**, *l*, and **p**. The set of parameter values used for the search are listed in Table 1. The algorithm for the optimization procedure is as follows,

1. Choose values for the robot link lengths: 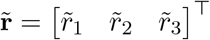.
2. Choose values for the human limb length and location: 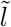 and 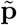.
3. Compute the set of all possible joint configurations for the human limb 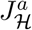 and the robot 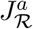 for the chosen robot and human limb parameters.
4. Compute the workspace 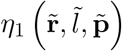 and robot manipulability measures 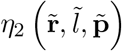.
5. If all possible human limb parameters have been searched, then go to Step 6, else go to Step 2.
6. Compute the two objective functions 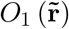 and 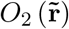 from the measures *η*_1_ and *η*_2_ computed for all possible human limb parameters.
7. If all possible robot parameters have been searched, then go to Step 8, else go to Step 1.
8. Find the optimal robot link parameters as the value of the **r** that maximizes *O* (**r**).

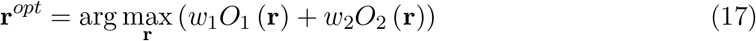 We assumed *w*_1_ = *w*_2_ = 0.5 for the current optimization problem.

**Table 1:**
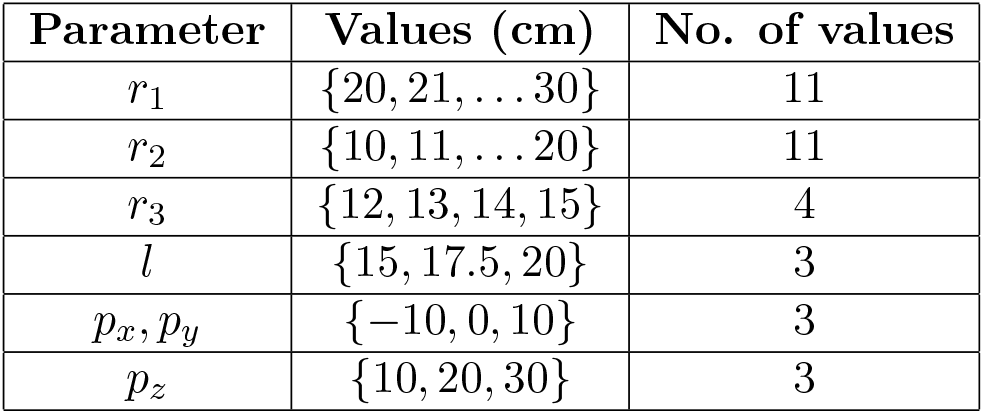
Set of parameter values for the robot and the human limb used for the robot link length optimization program. There are a total of 484 robot parameter sets (*r*_1_*, r*_2_*, r*_3_) that are searched, and for each of these 484 parameter sets the objective functions *O*_1_ (·) and *O*_2_ (·) are computed by averaging over the 8 different human limb parameters *K_l_* = 3*, K*_**p**_= 3.

#### 4.2 Simulation analysis of human limb parameter estimation

The algorithm for estimating the human limb parameters described in Section 3 was evaluated using simulated movement data. Twenty different random parameter sets were generated from uniform distributions for *p_x_*, *p_y_*, *p_z_*, and *l* of the human limb.

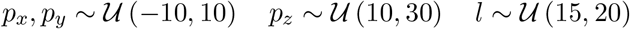

where, 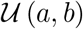 is a uniform probability density function with parameters *a* and *b*. The location and length of the human limb were sampled from the same region of parameters used for the optimization problem (Table 1). For each randomly selected parameter set **p** and *l*, 50 different random movements were imposed on the human-robot closed kinematic chain, and the human ****ϕ**** [*n*] and robot ****θ**** [*n*] joint configurations were recorded. The random movements to the human-robot closed chain were achieved by imposing a polysine movement to human joint of the following form,

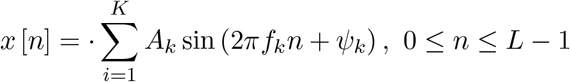

where, *K* is the number of sinusoidal components and was chosen as 3, *f_k_* = {0.2, 0.5, 1.0} *Hz*, *A_k_* = {1.0, 0.5, 0.1}, and *ψ_k_* was chosen randomly from a uniform distribution between −*π* and *π*. The polysine signal *x* [*n*] was appropriately scaled to cover a range of 0 deg to 90 deg for *ϕ*_1_, and −30 deg to 30 deg for *ϕ*_2_. The simulated data was assumed to be sampled at 100 Hz and 5 s of calibration movements were simulated. Gaussian white noise with two different variances 1 deg^2^ and 5 deg^2^ were added to the joint angles to simulate different levels of measurement noise; these two variances were chosen as these were considered to be reasonable noise variances for angles measured with a rotary encoder or a potentiometer. The robot parameters were assumed to be **r**^*opt*^ for these simulations.

The human and robot joint angle data from the simulated calibration procedures were used to estimate the human limb parameters; 50 different estimates for the 20 different sets of human parameters were estimated. The performance of the estimation algorithm was evaluated by computing the distribution of estimation errors for the four parameters.

## 5 Results

In this section, we present the results from the robot link length optimization and the simulation analysis of the human limb parameter identification algorithm.

### 5.1 Optimization of robot link lengths

The results from the robot link length optimization procedure are shown in Fig. 5 in the form of heatmaps as a function of two robot link parameters; the first two rows correspond to the individual objective functions *O*_1_ (**r**) and *O*_2_ (**r**), and the last row corresponds to the overall objective function *O* (**r**). The columns display these heatmaps as function of two of the robot link length parameters. As expected, the normalized workspace *O*_1_ (**r**) increases as the robot link lengths increase (first row of Fig. 5). On the other hand, the force ratio *O*_2_ (**r**) tends to be higher for shorter link lengths (Fig. 5). The overall objective function *O* (**r**), which is a weighted sum of *O*_1_ (**r**) and *O*_2_ (**r**) is shown in the third row in Fig. 5. Based on this plot and through numerical analysis, the optimal values for the robot link length parameters 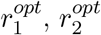, and 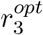 were found to be 27, 20, and 10 cm, respectively. Out of the 484 (11 × 11 × 4) robot link lengths searched, the set of robot link parameters that had the top 5% values for the objective function *O* (**r**) were found to have *r*1, *r*2, and *r*3 in the range 24 *−* 28 cm, 18 *−* 20 cm, and 10 cm, respectively. For the optimal link lengths, the range of values for of the normalized workspace *η*_1_ (**r**^*opt*^, *l*, **p**) and normalized force ratio *η*_2_ (**r**^*opt*^, *l*, **p**) for different values of the human limb parameters are depicted in Fig. 6. The top row shows the normalized workspace, which indicates that **r**^*opt*^, on average, is able to cover about 80% of the human limb’s workspace. The various values of the different parameters appear to result in a similar range of values for the normalized workspace, except for the *p_z_* = 10*cm* (rightmost figure in the top row in Fig. 6), where the normalized workspace is about 60%. Thus, when a subject sits very close to the robot, there is a drop in the subject’s workspace. The normalized force ratio (bottom row of Fig. 6) appear to be around 0.6-0.7, which means that in general 60-70% of the robot’s joint configurations have *ρ* (****θ****) ≥ 1. Based on the optimal link lengths, a 3D model of the proposed robot is depicted in Fig. 7, which shows two scenarios with the robot attached to the upper-arm and the forearm. When it is connected to the upper-arm, the robot can assist SFE and SAA. When it is attached to the forearm, it can support shoulder internal-external rotation and elbow flexion-extension, when the elbow position is constrained.

**Figure 5:**
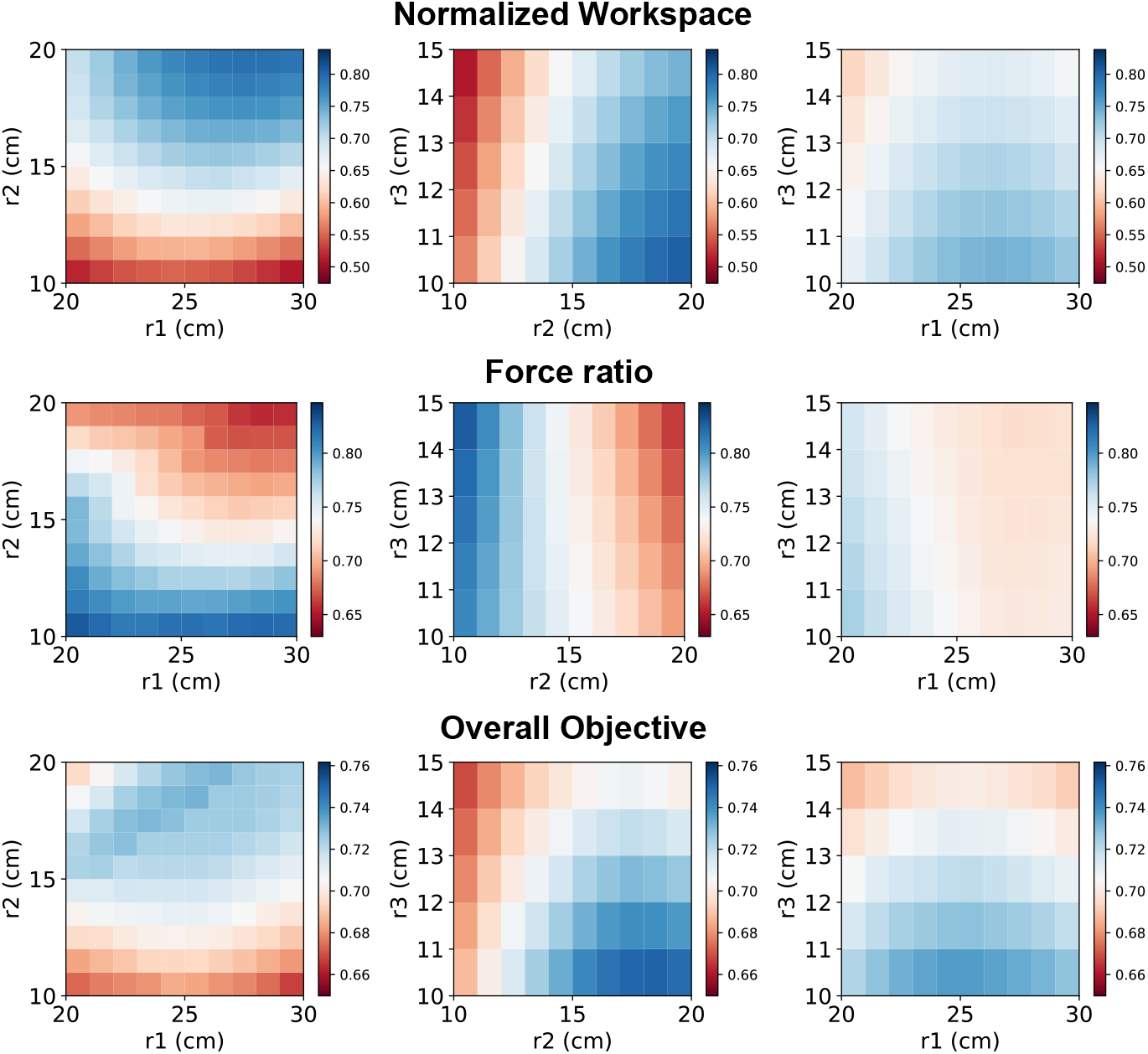
Heatmaps depicting the values of the objective functions *O*_1_, *O*_2_ and *O* as a function of the different robot link lengths. These plots show that workspace *O*_1_ and force ratio *O*_2_ are conflicting objectives, and the resulting overall objective that weighs both *O*_1_ and *O*_2_ equally is shown in the bottom row.

**Figure 6:**
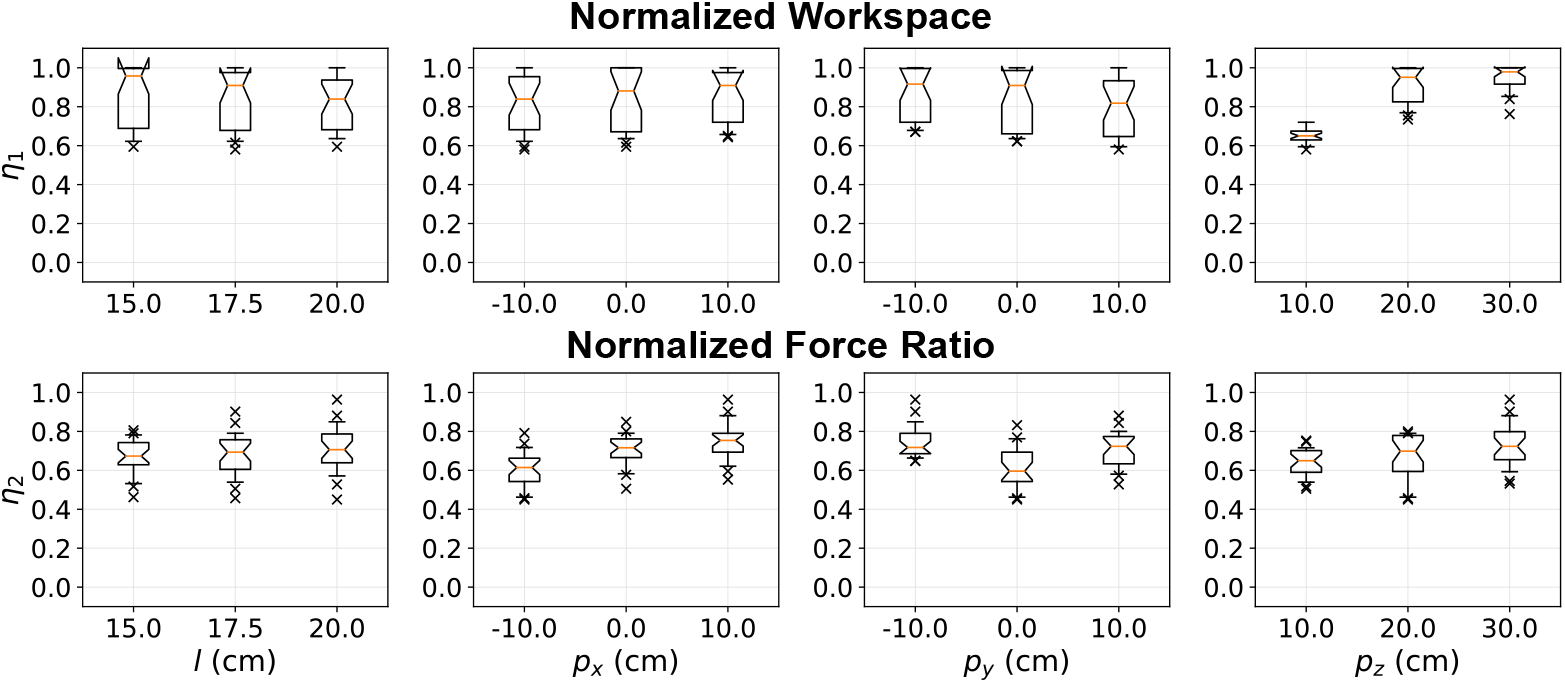
The values of normalized workspace *O*_1_ and force ratio *O*_2_ for the optimal robot link lengths, for the different possible parameters for the human limb (from Table 1).

### 5.2 Identification of human limb parameters

The results from the analysis of the human limb parameter identification procedure are shown in Fig. 8. The true value of the parameters is **p**, the estimated parameter is 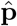, and estimation error ****ϵ****, its mean ****μ_ϵ_**** and covariance **C**_*ϵ*_ are given by,

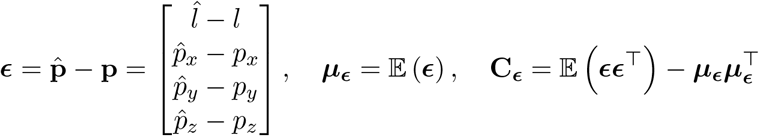

where, 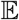 is the expectation operator.

**Figure 8:**
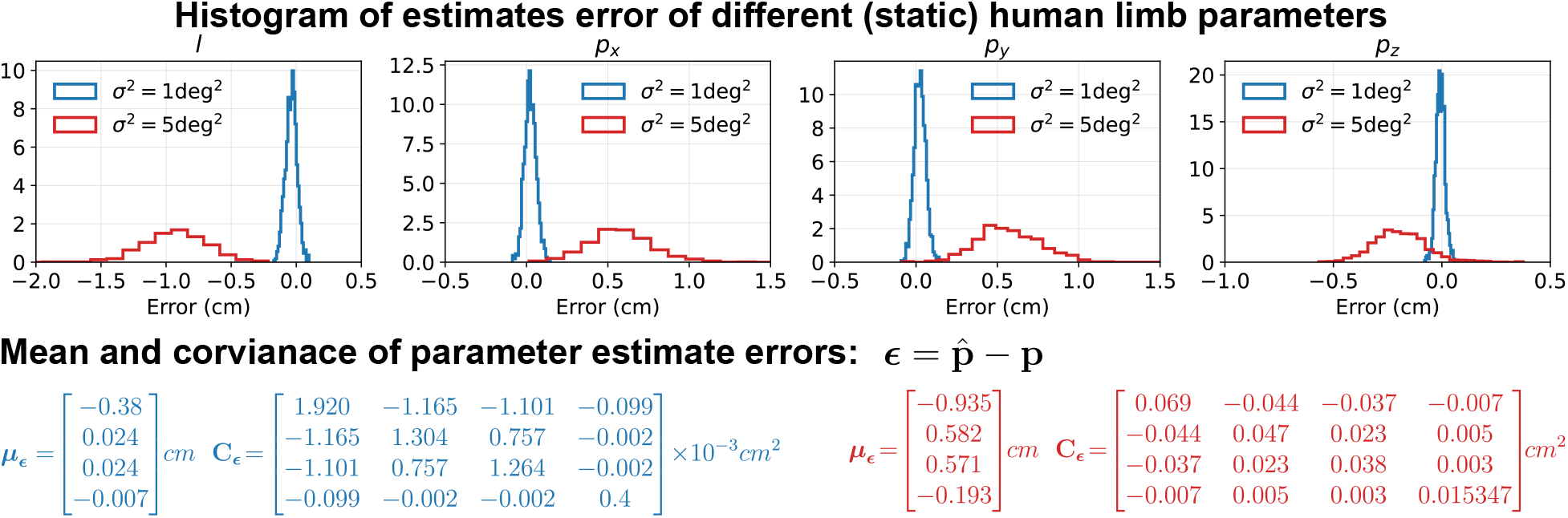
Errors in human limb parameter estimates for two different measurement noise *σ*^2^ = 1 deg^2^ and *σ*^2^ = 5 deg^2^. The mean and the covariance of the different parameter estimates are shown below the histograms. The text in blue color represent to *σ*^2^ = 1 deg^2^, while the one in red corresponds to *σ*^2^ = 5 deg^2^.

Fig. 8 shows histograms of the estimation error in the different parameter estimates for two different noise levels in the joint angle measurements. As expected, lower measurement noise results in estimates with smaller bias and variance. The sample mean and covariance of estimation errors are shown below the plots in Fig. 8 for both noise variances. The limb length has the largest absolute bias and variance among the four parameters, and also appears to be underestimated for both levels of measurement noise. The parameter *p_z_* has the lowest absolute bias and variance, which also appears to be underestimated. The other two parameters *p_x_* and *p_y_* have intermediate bias and variances, and these are overestimated. *p_z_* is least correlated to the other parameters. The other three parameters appear to be slightly correlated.

## 6 Discussion

The paper presented the kinematic design of a 6-*dof* robot - AREBO - capable of assisting movements of individual human joints. The proposed design can assist up to two *dof* of the human joint while ensuring the safe application of assistive forces on the human limb. The optimization of the robot’s link lengths, maximizing an objective function consisting of the human limb’s workspace and the robot’s ability to apply forces in safe directions, was presented. The paper also presented a simple algorithm for continuously estimating the kinematic parameters of the human limb using the joint angles of the robot and the human limb.

Based on the existing evidence and further assuming that robot-assisted therapy is as effective as dose-matched conventional therapy, the ultimate goal of rehabilitation robots is to deliver substantial doses of intense therapy at a small cost to the healthcare system. Realizing this goal requires compact, cost-effective devices that can be easily deployed even in space-constrained healthcare settings and patients’ homes while offering superior benefit-to-cost ratio to the user. AREBO presents a minimalistic solution for an arm robot by using three actuated and three passive *dof*, while offering several useful features that boost its potential for clinical adoption.

1. AREBO has a very compact and portable structure making it suitable for small clinics and even patients’ homes.
2. The reduced number of actuators also makes the overall bill of materials low compared to that of a fully actuated robot.
3. From the perspective of assisting just two *dof* of the human limb, AREBO’s self-aligning passive joints remove the need for active alignment, thus simplifying the control of the human-robot interaction.
4. The end-effector design of the robot’s structure allows it to be used for both the left and right arms without any change in its structure. Without such a feature, a clinic would require dedicated devices for the left and right arms, which is an undesirable solution.
5. The robot can be used for training one or two *dof* at either the shoulder or the elbow, as shown in Fig. 7. SFE and SAA can be trained with the design shown in Fig. 7(a), while the shoulder internal-external rotation *dof* can be trained by connecting the robot to the forearm (Fig. 7(b)), which can also be coupled with the elbow flexion-extension. It would be safer and more comfortable to assisted shoulder internal-external rotation by applied forces on the forearm [15].

Another important feature of AREBO is the relaxed constraint on the patient’s relative position with respect to the robot. This feature is of significant practical value, as this has the potential to drastically reduce setup time for patients with more severe impairments or in a wheelchair. This ease in setting up the device translates to improved usability and can save time for the clinician when using the robot with multiple patients during a day. This reduced constraint in seating the patient and variations in human limb size between patients result in variations in the human limb’s workspace that can be supported by the robot, and also its manipulability. However, as long as these parameters are within a reasonable range specified in Table 1, the robot with optimized link lengths has an excellent workspace and a good manipulability. The optimized robot can cover, on average, 80% of the human limb’s workspace, and for about 60-70% of the points in the robot’s joint space, it is “easier” to apply a force orthogonal to the human limb.

The proposed algorithm for human limb parameter estimation allows AREBO to automatically estimate the location of the human joint with respect to the robot, and the length of the human limb, both of which are required for the complete specification of the human-robot closed kinematic chain. This can be done with a short calibration procedure (5 s long). The current results show that with the level of noise expected from rotary sensors, the human limb parameters can be estimated with relatively small bias and variance. There has been prior work on estimating human limb posture when connected to an exoskeleton robot [16], and to plan the robot’s trajectory for a given human limb trajectory [15]. However, the authors are unaware of any prior work on estimating the location of the human joint and limb length using the human and robot joint kinematic data. One of the assumptions made by the proposed algorithm is that the human limb parameters are fixed over time. It is reasonable to assume that limb length does not change over time, but the same cannot be assumed about the limb’s location. For example, when the robot is connected to the upper-arm (as shown in Fig. 7(a)), then trunk movements or scapular movements will result in the translation of the glenohumeral joint. Thus, it would be ideal if the estimation can be implemented using data from the recent past in a recursive form with the following assumptions:

1. Human limb length does not undergo any change over time.
2. The location of the human limb does not undergo a drastic change in the window of data used for the estimation process.

Based on these assumptions, the recursive estimate at time *n* can be posed as multiple objective minimization problem,

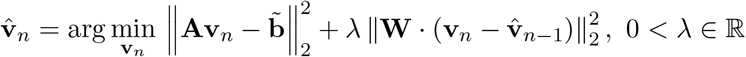

where, 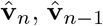 are the parameter estimated at time *n* and *n −* 1, respectively. **W** is a weight matrix that assigns different weights for change in different parameters (e.g., the weight for a change in *l* can be set much higher than the other parameters). The ability to track changes in human limb parameters can allow the robot to automatically detect compensatory movements within and across sessions, which can be a useful measure of motor ability [17].

One potential limitation of AREBO is that it cannot be used for the assisted training of multi-joint movements possible with exoskeleton robots such as BONES [10], ArmIn [8], etc. From the neurorehabilitation perspective, the current evidence indicates that simple single-joint training is as effective as complex multi-joint training [4]. The study by Milot et al. showed that both single and multi-joint training with BONES resulted in improvements in both sensorimotor impairments and function [4]. About 75% of the participants in this study had an equal preference for both single-joint and multi-joint training with the robot [4]. There is also evidence for the transfer of training effects to untrained ADL activities [5], which can be extrapolated to the possibility of single-joint training generalizing to complex multi-joint movements. Furthermore, single-joint exercises are an important component of training protocols in patients with severe sensorimotor impairments in impairment oriented training [18, 19]. These studies and the practical advantages of AREBO make a strong case for developing and evaluating the clinical usability and efficacy of such simple, compact robots for arm neurorehabilitation. If found to be therapeutically effective, such robots have the best potential for clinical translation and widespread adoption.

## 7 Conclusion

The paper presented the kinematic design and optimization of a compact 6-*dof* robot for individual joint training of the human arm. The proposed robot uses three actuated *dof* and three passive self-aligning *dof* keeping the overall structure of the robot simple and cost-effective. The proposed robot allows significant freedom in terms of the seating of a patient with respect to the robot, can easily accommodate arms of different sizes, and can be used for both the left and right arms without any change to its structure. The paper also presented an approach for automatically tracking the kinematic parameters of a human limb attached to the robot, which can be used by a robot controller to impose the desired movements to the human limb.

## Contributorship

SB, SG, and SS conceived the initial ideas for the robot and its design. SG and SB explored various potential designs for the robot. SB carried out the optimization of the robot link lengths. SB, SG, and JSM worked on the estimation algorithm and its evaluation through simulated data. SB prepared the first draft of the manuscript with help from SG and JSM. All authors reviewed the manuscript. SB and SS oversaw the activities of the project.

## Acknowledgements

The authors thank Mr. Aravind Nehrujee, Mr. Samuel Elias, and Mr. Sathish Balaraman for their help in building and rendering the 3D models for the robot.

## Appendix A: Inverse Kinematics of AREBO

Given a desired endpoint position **p**_*d*_ and orientation **R**_*d*_ for the robot’s endpoint, we find out if the robot can reach this position and orientation, and if so we would like to compute the ****θ**** that would help achieve this endpoint position and orientation.

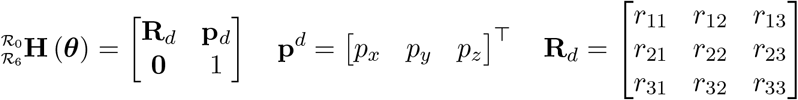

The angle *θ*_1_ can be determined from the *x* and *y* components of **p**_*d*_ (Eq. 2).

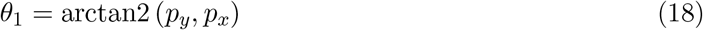

The inverse position kinematics sub-problem can now be converted to a 3-link planar arm problem by substituting the value of *θ*_1_; the three links are *r*_1_, *r*_2_, and *r*_3_ in Fig. 3. We first compute two quantities, *x_d_* and *y_d_* as the following, from Eq. 2,

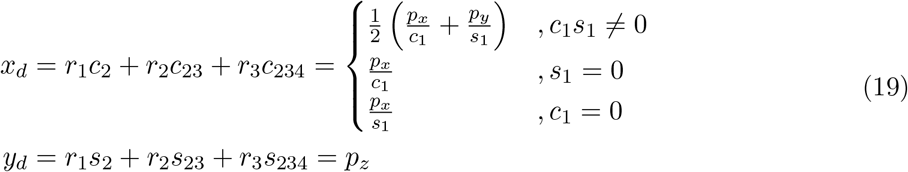

The inverse kinematics of the 3-link planar arm does not have a unique solution. However, the angle of the third link (*c*) in the 3-link planar arm, i.e., *θ*_2_ + *θ*_3_ + *θ*_4_. This can be obtained by solving the inverse orientation kinematics problem.

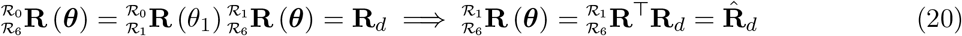

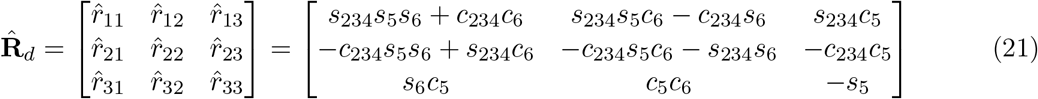

We can now solve for 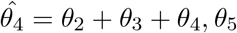 and *θ*_6_ using the following,

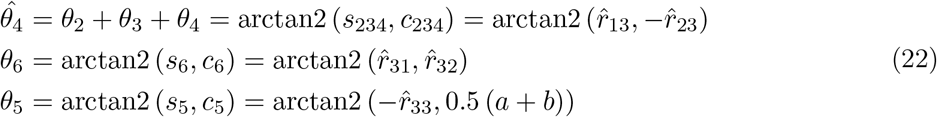

where,

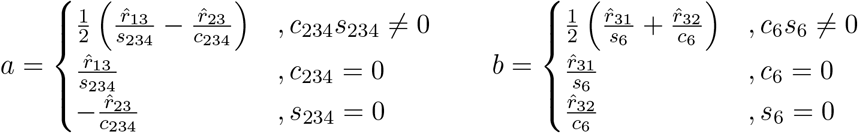

Next, 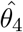 can be used to solve the 3-link planar arm inverse position kinematics,

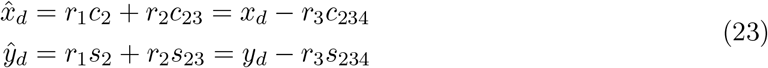

We can solve for *θ*_2_ and *θ*_3_ using the standard inverse kinematics procedure for a 2-link planar arm using 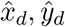.

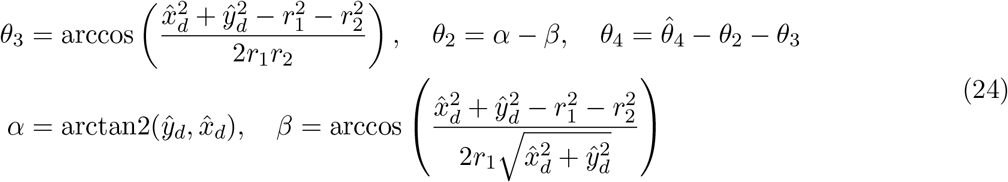

## Appendix B Analysis code

The code for all the analyses presented in the manuscript is available for downloaded from here.

## References

[1] GB Prange, MJA Jannink, CGM Groothuis-Oudshoorn, HJ Hermens, and MJ Ijzerman. Systematic review of the effect of robot-aided therapy on recovery of the hemiparetic arm after stroke. Journal of rehabilitation research and development, 43(2):171–184, 2009.

[2] Gert Kwakkel, Boudewijn J Kollen, and Hermano I Krebs. Effects of robot-assisted therapy on upper limb recovery after stroke: a systematic review. Neurorehabilitation and neural repair, 22(2):111–121, 2008.

[3] Albert C Lo, Peter D Guarino, Lorie G Richards, Jodie K Haselkorn, George F Wittenberg, Daniel G Federman, Robert J Ringer, Todd H Wagner, Hermano I Krebs, Bruce T Volpe, et al. Robot-assisted therapy for long-term upper-limb impairment after stroke. New England Journal of Medicine, 362(19):1772–1783, 2010.

[4] Marie-Hélène Milot, Steven J Spencer, Vicky Chan, James P Allington, Julius Klein, Cathy Chou, James E Bobrow, Steven C Cramer, and David J Reinkensmeyer. A crossover pilot study evaluating the functional outcomes of two different types of robotic movement training in chronic stroke survivors using the arm exoskeleton bones. Journal of neuroengineering and rehabilitation, 10(1):112, 2013.

[5] Sydney Y Schaefer, Chavelle B Patterson, and Catherine E Lang. Transfer of training between distinct motor tasks after stroke: implications for task-specific approaches to upper-extremity neurorehabilitation. Neurorehabilitation and neural repair, 27(7):602–612, 2013.

[6] Gerard G Fluet, Alma S Merians, Qinyin Qiu, Amy Davidow, and Sergei V Adamovich. Comparing integrated training of the hand and arm with isolated training of the same effectors in persons with stroke using haptically rendered virtual environments, a randomized clinical trial. Journal of neuroengineering and rehabilitation, 11(1):126, 2014.

[7] Julius Klein, Steven J Spencer, and David J Reinkensmeyer. Breaking it down is better: haptic decomposition of complex movements aids in robot-assisted motor learning. IEEE Transactions on Neural Systems and Rehabilitation Engineering, 20(3):268–275, 2012.

[8] Tobias Nef, Marco Guidali, and Robert Riener. Armin iii–arm therapy exoskeleton with an ergonomic shoulder actuation. Applied Bionics and Biomechanics, 6(2):127–142, 2009.

[9] RJ Sanchez, Eric Wolbrecht, R Smith, J Liu, S Rao, S Cramer, T Rahman, James E Bobrow, and David J Reinkensmeyer. A pneumatic robot for re-training arm movement after stroke: Rationale and mechanical design. In 9th International Conference on Rehabilitation Robotics, 2005. ICORR 2005., pages 500–504. IEEE, 2005.

[10] J Klein, SJ Spencer, J Allington, K Minakata, ET Wolbrecht, R Smith, JE Bobrow, and DJ Reinkensmeyer. Biomimetic orthosis for the neurorehabilitation of the elbow and shoulder (bones). In 2008 2nd IEEE RAS & EMBS International Conference on Biomedical Robotics and Biomechatronics, pages 535–541. IEEE, 2008.

[11] H Igo Krebs, Neville Hogan, Mindy L Aisen, and Bruce T Volpe. Robot-aided neurorehabilitation. IEEE transactions on rehabilitation engineering, 6(1):75–87, 1998.

[12] Domenico Campolo, Paolo Tommasino, Kumudu Gamage, Julius Klein, Charmayne ML Hughes, and Lorenzo Masia. H-man: A planar, h-shape cabled differential robotic manipulandum for experiments on human motor control. Journal of neuroscience methods, 235:285–297, 2014.

[13] Che Fai Yeong, Alejandro Melendez-Calderon, Roger Gassert, and Etienne Burdet. Reachman: a personal robot to train reaching and manipulation. In 2009 IEEE/RSJ International Conference on Intelligent Robots and Systems, pages 4080–4085. IEEE, 2009.

[14] Giulio Rosati, Paolo Gallina, and Stefano Masiero. Design, implementation and clinical tests of a wire-based robot for neurorehabilitation. IEEE Transactions on Neural Systems and Rehabilitation Engineering, 15(4):560–569, 2007.

[15] Nathanaël Jarrassé and Guillaume Morel. Connecting a human limb to an exoskeleton. IEEE Transactions on Robotics, 28(3):697–709, 2011.

[16] Camilo Cortés, Aitor Ardanza, F Molina-Rueda, A Cuesta-Gómez, Luis Unzueta, Gorka Epelde, Oscar E Ruiz, Alessandro De Mauro, and Julian Florez. Upper limb posture estimation in robotic and virtual reality-based rehabilitation. BioMed research international, 2014.

[17] MC Cirstea and Mindy F Levin. Compensatory strategies for reaching in stroke. Brain, 123(5):940–953, 2000.

[18] T Platz. Impairment-oriented training (iot)–scientific concept and evidence-based treatment strategies. Restorative neurology and neuroscience, 22(3-5):301–315, 2004.

[19] Thomas Platz, Stefanie van Kaick, Jan Mehrholz, Ottmar Leidner, Christel Eickhof, and Marcus Pohl. Best conventional therapy versus modular impairment-oriented training for arm paresis after stroke: a single-blind, multicenter randomized controlled trial. Neurorehabilitation and neural repair, 23(7):706–716, 2009.

